# Wildlife and Marine Mammal Spatial Observatory: Observation and automated detection of Southern Right Whales in multispectral satellite imagery

**DOI:** 10.1101/2022.01.20.477141

**Authors:** Ludwig Houegnigan, Enrique Romero Merino, Els Vermeulen, Jessica Block, Pooyan Safari, Francesc Moreno-Noguer, Climent Nadeu

## Abstract

The Wildlife and Marine Mammal Spatial Observatory is a joint research effort for the census of *wildlife and particularly of marine mammals in satellite imagery. In that context, this paper illustrates the development of a high accuracy algorithm for the detection of right whales in sub-meter resolution multispectral satellite imagery with the constraint of a relatively small sample support of 580 southern right whale images. A significant space is devoted to exploratory data analysis to describe the statistical structure of right whale pixels and ocean surface pixels across multispectral bands.*

*Observations of southern right whale in satellite imagery are divided into typical and atypical right whale forms and the first observations of right whale mother and calf pairs in satellite imagery are presented. Measurements of whales are furthermore automatically extracted from whale observations (major axis length, minor axis length, etc). A significant space is also devoted to statistical data exploration, a step frequently overlooked in machine learning solutions, that yet offers interesting insight into the structure of animal detection in satellite imagery. The extracted statistics can readily be used by researchers to develop detection solutions even with low sample support. The adopted solution for detection consists of feature extraction with a convolutional neural network followed by classification with a support vector machine. 20 different convolutional neural networks were tested for feature extraction. Biostatistics parameters (accuracy, sensitivity, specificity and precision) were measured for comparison. Most architectures generally achieved high performance with low false positive and false negative rates. 100% accuracy is achieved in the case of 2 convolutional neural networks, Nasnet Large and Inception V3, and only with a specific selection of multispectral bands.*

NB: This is a preprint that does not include satellite imagery due recent reviews

## Introduction

### 1.1 Context of the Wildlife and Marine Mammal Spatial Observatory (WIMMSO)

The Wildlife and Marine Mammal Spatial Observatory is an effort led primarily by UPC BarcelonaTech with contributions from partners at UC San Diego and further collaboration with researchers engineers, data scientists, marine mammal scientists, oceanographers and conservationists from several institutions worldwide. It proposes to analyze satellite imagery in order to help researchers and conservationists to census, answer research questions and acquire more knowledge on the habitat, migration routes, feeding and breeding grounds of endangered and data-deficient species. A key step in this endeavor is the development and analysis of a database of observations per species that is an implicit prerequisite for the development of accurate detection, classification and localization algorithms of megafauna in satellite imagery.

A first description of the Wildlife and Marine Mammal Spatial Observatory through the Global Scan and Pacific Scan projects was introduced at the World Marine Mammal conference [1], from December 9-12, 2019, in Barcelona, Spain. Enhanced image visualizations, early results for visual and automated detections for multiple species (right whales, gray whales, beluga whales, humpback whales) as well as an early count of the detected animals per species were presented with an actualized count of the number of animals. The idea of gathering global sightings for a large number of species by surveying in priority their typical aggregations was introduced. Species were identified by context (reported sightings, seasonal aggregation, likelihood and physical resemblance) while others had to remain unassigned.

In this paper, we investigate deeper into a particular species, namely right whales of the genus *Eubalaena.* A specific statistical analysis is produced based on a large and, to the best of our knowledge, unprecedented number of individual animals of this species, as well as image enhancements and tools to facilitate both the visual and automated detection of right whales in submeter resolution satellite imagery.

In this particular case, the ability to consistently monitor the dynamics of fragile right whales populations through satellite imagery can be a key asset and to acquire knowledge on distribution and migration to support their conservation. Furthermore, the abundance of observation of southern right whales can be used as a proxy for the detection of the less abundant and critically-endangered Northern Right Whales and North Pacific Right Whales. This paper furthermore focuses on the observation of right whales in satellite imagery in South African coastal waters. We observe them through multiple sensors and multiple postures, and include observation of mother and calf pairs, which to the best of our knowledge was never documented before.

For the sake of concision, it is beyond the scope of this paper to build an optimal detector or to delve into detection theory, which would require its own detailed publication. Rather, the key points of this paper are firstly to present image processing tools that are helpful for the analysis and visualization of large and complex remote sensing datasets, secondly to put more focus on a statistical and descriptive analysis of the satellite images, a step which is frequently overlooked. In the context of satellite images that can be rather limited and costly, the extracted statistical knowledge presented here is relatively easily transferable for readers who do not have access to a sufficiently large database of images, in order to build their own detectors and classifiers. Last but not least, this is also an opportunity to acknowledge and make space for the technical and visual quality of the observation of wildlife in satellite imagery.

### 1.2 Species and areas of interest

#### 1.2.1 A focus on Right Whales

The Eubalaena genus is constituted by three distinct phylogenetic species: Southern Right Whales (Eubalaena australis), North Atlantic Right Whales (Eubalaena glacialis), and North Pacific Right whales (Eubalaena japonica) that have spatially non-overlapping distributions. The genus faced the pressure of intensive whaling until its international protection in 1935. Even though recovery from past threats has been ongoing for decades for southern right whales [2,3,4,5], populations remain in a precarious state and vulnerable to threats of more recent concern such as climate change and its impact on their food availability [6,7]. The recovery of North Atlantic right whales has not been so successful, entanglement in fishing nets as well as ship strikes have proven to have a major impact on populations [8]. Furthermore, events such as Unusual Mortality Events (UME) that are difficult to predict can impose a heavy toll on populations at risk: an unusual Mortality event initiated in 2017 [9] has seen the death of 45 individuals which is a significant impact for the critically-endangered population of North Atlantic right whales which numbers approximately 400 individuals is declining [10] and exhibits poor body conditions [11]. Such unpredictable events generally highlight the precariousness of their status even when positive trends of population growth are observed for certain species.

A 2018 study based on passive acoustic monitoring [12] pointed to a reduced acoustic activity of North Atlantic right whales in the northern Gulf of Maine, which in turn could be hypothesized either as a significant decline in population size or as a change in spatial distribution. The latter was later confirmed by efforts performed in Canada [13] indicating an increased vocal activity of North Atlantic right whales in the Gulf of Saint Lawrence, Canada. This points to the need of thorough and reactive spatial and temporal monitoring, that could possibly be provided by monitoring based on satellite imagery, for species with precarious status and which behavior may be rapidly altered by a changing climate [14].

The phylogenetic relationship as well as morphological similarities between the different populations of right whales allows to study the Eubalaena genus as a whole from the point of view of image analysis and processing [11]. Even though the critical status of North Atlantic and North Pacific right Whales lends itself more to intensive research and conservation effort, in the context of the Wildlife Marine Mammal Spatial Observatory that relies on computer vision and machine learning techniques, building a relatively large database of observations will improve our ability to detect whales in satellite imagery, and thus, efforts were focused on the large winter congregations of Southern Right Whale. Furthermore, gathering a large database of observations obtained in large aggregations will allow us to better study the underlying statistics for the problem of detection of right whales in satellite imagery, a task which has not been carried out so far since most work so far relies on a relatively small observational sample support. The robust knowledge acquired for abundant populations can then easily transfer to the more endangered populations.

While a benchmark paper studied the presence of right whales in satellite imagery in Peninsula Valdes, Argentina [67], a focus is here given to the presence of Southern Right whales in South Africa for reasons which will become explicit in section 1.2.3.

#### 1.2.2 Southern Right Whales and the South African population

Whereas the populations of North Atlantic right whales and North Pacific Right whales are endangered and *number very few animals,* the global southern right whale population was estimated in 2009 at 13,600 individuals [15] distributed across the South Atlantic South Pacific and Indian oceans. These whales perform yearly migrations from wintering calving sites to summering sub-Antarctic waters where they feed on large aggregations of Antarctic Krill *(Euphausia superba)* and copepods. The largest winter aggregations occur in Peninsula Valdes (Argentina) [16,17], southern Cape coast (South Africa) [18], Australia particularly between Head of Bight and Fowlers Bay [19], as well as in New Zealand [20, 21, 22], Brazil [23, 24, 25] and Uruguay [26, 27]. In smaller numbers sightings have been reported of the coasts off Peru and Chile [28, 29, 30] where photo-identification [31]and UAV-based fieldwork led to the hypothesis of the existence of a nursery ground [32].

However, in recent years, fluctuations in Southern Right Whale coastal prevalence across these wintering grounds population trends have been observed with elongation of calving intervals [33,34,35], and short-term (at least) effects on population growth rate [36]. These distribution shifts are more extreme in South Africa with sightings of cow-calf pairs decreasing under 80% of the norm [24].

Even though the estimated number of whales wintering in South Africa is considered to be less than in Peninsula Valdes, the South African breeding ground has characteristics of particular interest for surveys based on satellite imagery: water shallowness and low turbidity in that coastal nursery has allowed us to unequivocally observe a large number of Southern Right whales from head to tail in the imagery (cf. section 2.2.5). The area furthermore offers favorable conditions for right whales to rear their calves [37] while allowing us to carry out meaningful observation and presence estimation of the animals. As a result, the presence of calves and juveniles with smaller size and dark to fair skin tones is clearly observed and documented here, which to the best of our knowledge is an unprecedented observation in satellite imagery.

#### 1.2.3 Focus on South Africa and description of the study area

The present study includes satellite imagery over the Agulhas ecoregion in the warm and temperate South Coast of South Africa which is itself a part of the Greater Cape Floristic Region of South Africa, a biodiversity hotspot where endemism and species richness are among the highest globally while facing significant anthropogenic pressure and being rapidly impacted by climate change. The Agulhas ecoregion features a warm temperate climate due to the mixing of waters from south-western Africa via the colder Benguela current, with subtropical and sub-Antarctic waters via the warmer Agulhas current [46,47]. Such mixing zones are generally known for their higher productivity which can support a large number of species. As a consequence, the region indeed has the highest number species within South Africa and is a breeding area for many of them. The area is characterized by rocky shores, sand shores and includes biodiversity hotspots such as Hermanus’ whale sanctuary in Walker Bay or the De Hoop Nature Reserve, a UNESCO World Heritage Site since 2004. The latter includes the De Hoop Marine Protected Area, declared in 1985 which covers 51km of protected shoreline and an area of approximately 253 km2. The De Hoop MPA provides habitat for dolphins, seals, southern right whales and at least 250 species of fish many of which are endemic. As a no-catch area, the MPA has provided refuge for several overexploited species and has demonstrated important benefits for the protection of economically important inshore reef fish species such as galjoen, wildeperd, blacktail, black musselcracker, and white musselcracker. It generally provides an area where anthropogenic impact is reduced which has demonstrably been beneficial to a large number of species.

The southern right whale population that utilizes the nursery ground off the southern Cape coast of South Africa has been studied intensively since the early 1970s through a series of annual aerial surveys between Nature’s Valley and Muizenberg, in order to identify individuals using photography of natural markings [38–43].

The distribution of southern right whales along the southern Cape coast breeding ground is non-random, with different sections of the coast utilized as preferential habitat by these two components of the population. Reason for this non-uniform distribution has been ascribed to habitat selection of nursing cows. In fact, the majority of cow-calf pairs seem to concentrate in areas that provide protection from swell and winds as well as rocky bottoms. Shallow water, sandy bottoms and gentle slopes are preferred for an improved energy conservation for lactating females and protection against possible mating attempts of adult males and against predation risk on new-born calves [28].

**Figure 1.1.**
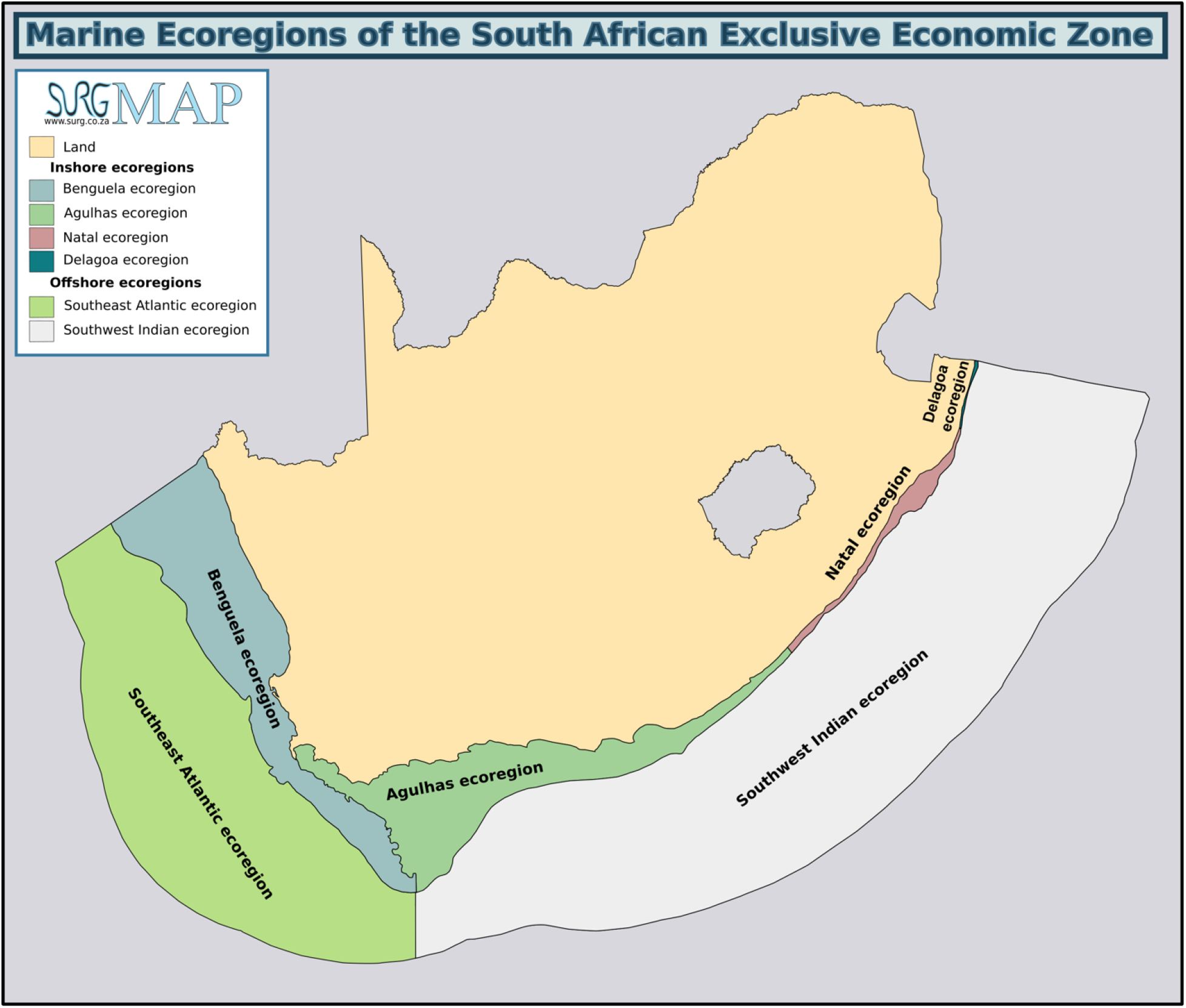
Marine Ecoregions of the South African Exclusive Economic Zone, used under CMM licence.

Southern right whale aggregations occurring in the Agulhas ecoregion feature a large number of cow-calf pairs and seem to offer an interesting situation for visualization with moderate cloud coverage, low water turbidity, calm bays and a strong coastal distribution. The site of Hermanus and De Hoop Nature Reserve are certainly the most renowned and accessible for whale observation but aggregations typically can occur in a series of bays all the way between False Bay and Plettenberg Bay. Southern right whale presence is evidenced from June to December with a peak in late September, a time by which most calves are considered to be already born [43]. Annual aerial surveys therefore typically take place in early October [44,45].

## 2. Observation of Southern Right Whales through multiple sensors

**Fig.2.1.** Example of observation of southern right whales through satellite imagery. A small aggregation of Southern Right Whales can be observed near the De Hoop Reserve. Low turbidity and a sandy and light reflective bottom allow to spot the southern right whales clearly in their calving ground.

**Fig 2.2.** Satellite image with 4 southern right whales. Image enhancement while not conserving natural colors makes body color nuances as well as potential water splashes appear more clearly on the whale bodies.

### 2.1 Data description

52 satellite images spanning a time period from 2002 to 2019 were gathered and analyzed. Over this period of time, multiple sensors have been operational and were included among which Ikonos, GeoEye, World View 1, World View 2, and World View 3. While the different multispectral bands have standardized names (blue, red, green, panchromatic, near infra-red, etc), this nomenclature poorly reflects the variability in data acquisition, spectral response and the different filter shapes characterizing each band which inevitably translates in visual and numerical differences in the acquired images. Among the sensors available for this study the number of bands varies from 1 to 8, yet, even for bands that have the same denomination (panchromatic, green, red blue) significant differences exist across sensors. For example, the panchromatic band in GeoEye and Worldview 2 are respectively 450-800nm and 400-900nm, a 30% relative bandwidth difference, while the near infrared band in Quickbird and Ikonos featured a 31% relative bandwidth difference (Table 2.1). These aspects as well as notably different spectral responses can result in offsets across nominally similar bands.

**Table 2.1.** Characteristics of available sensor data.

|  | Ikonos | Quickbird | GeoEye | WVo1 | WVo2 | WVo3 |
| --- | --- | --- | --- | --- | --- | --- |
| Number of images |  |  |  |  |  |  |
| Years active | 1999-2015 | 2001-present | 2008-present | 2007-present | 2009-present | 2013-present |
| Panchromatic<br>GSD (cm) at<br>nadir/GSD at<br>20deg off<br>nadir PAN | 82/100 | 65/73 | 0.46/NC | 50/55 | 46/52 | 31/34 |
| Multispectral<br>GSD at nadir<br>(cm)/GSD at<br>20deg off<br>nadir(cm) | 328/400 | 262/290 | 184 /NC | - | 184 /240 | 124/138 |
| Frequency<br>Bands | 4MS +1<br>PAN | 4MS +1<br>PAN | 4MS +1<br>PAN | 1PAN | 8 MS+1<br>PAN | 8MS +1PAN |
| Coastal range | - | - | - | - | 396-458nm | 400 - 450<br>nm |
| Blue Range | 0.445- 0.516<br>um | 450-520 | 450 - 510 | - | 442-515 | 450 - 510 nm |
| Green Range | 0.506<br>-0.595um | 520-600 | 510 - 580 | - | 506-586 | 510 - 580 nm |
| Yellow | - | - | - | - | 584-632nm | 585 - 625 nm |
| Red | 0.632-0.698um | 630-690 | 655 - 690 | - | 630 - 690<br>nm | 630 - 690<br>nm |
| Red-Edge | - | - | - | - | 699-749 nm | 705 - 745 nm |
| NIR<sub>1</sub> | 0.757<br>-0.853um | 760-900 | 780 - 920 | - | 765-901 nm | 770 - 895 nm |
| NIR<sub>2</sub> | - | - | - | - | 856-1043nm | 860 - 1040<br>nm |
| Panchromatic | 0.45-0.90 | 450-900 nm | 450 - 800 | 400-900nm | 400-900 nm | 450-800 nm |

As technology improved through time, ground sampling distance at nadir and off-nadir also varies greatly from a sensor to another and such diversity should be taken into account in the development of detection and classification methods. Given this variability and for the sake of consistency, the statistical data analysis presented in section 3 focuses mainly on 580 detections of Southern Right Whale from the Worldview 3 sensor. For the sake of concision, additional results presenting the detection performance in images captured with other sensors and generalization to other areas will be presented in a later publication.

### 2.2 Feature Extraction using pretrained networks and Support Vector Machines

Feature extraction is a convenient manner to use the performance and discriminative capability of deep neural networks pretrained on larger datasets with more than a million images such as Imagenet or Places 365, and to transfer those abilities to a less well-defined problem with a smaller sample support such as the current right whale detection problem. Using one of the final layers of a deep neural network, robust and general image features can be extracted. Then a support vector machine (SVM) can be trained on labelled extracted features with results that are typically better than training from scratch or than fine-tuning a deep neural network when the small sample support for training is small. As an example, feature extraction with the deep neural network Alexnet is illustrated in figure 2.3, for which activations on the fully connected layer ‘fc6’ are used to extract adequate features which are then fed to an SVM.

**Figure 2.3.**
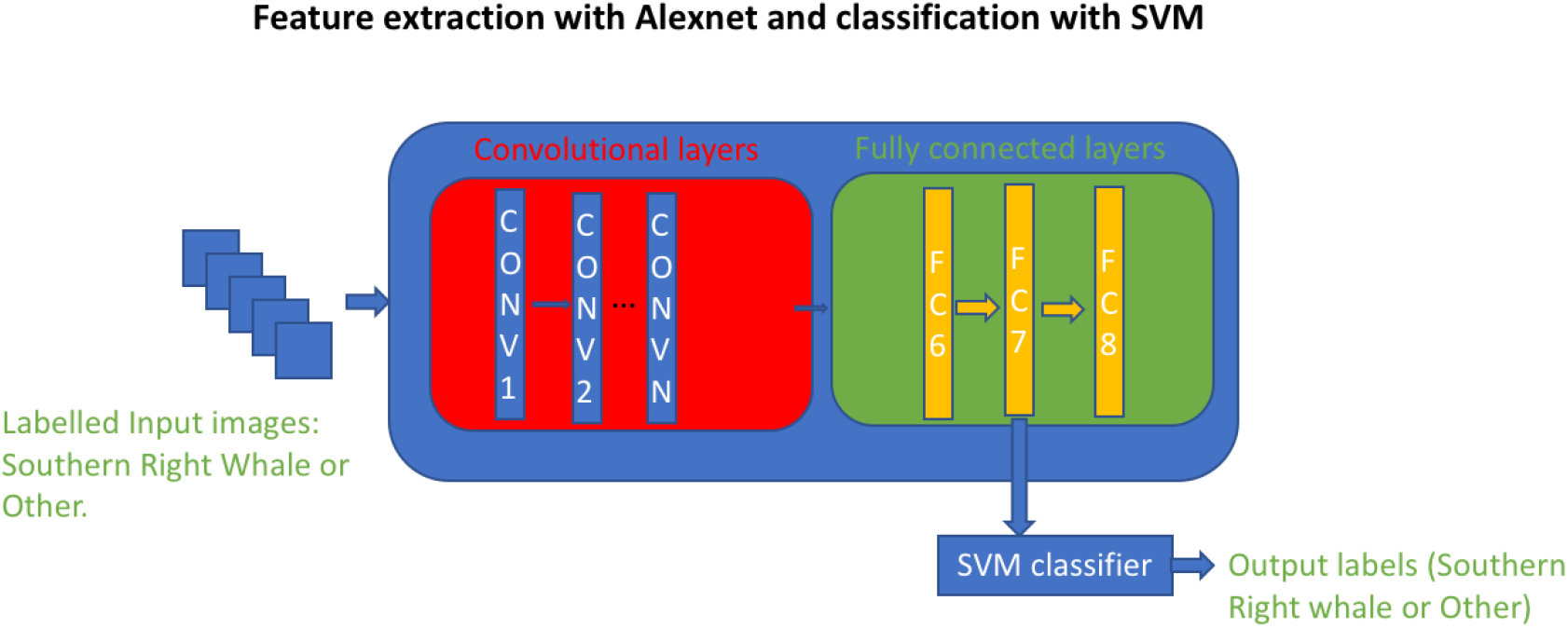
Feature Extraction with Alexnet.

While Alexnet was at the origin of a significant breakthrough for image recognition, significant progress has been made for the task and multiple deep neural networks have since outperformed Alexnet [46]. For the purpose of our recognition task, the following 20 different deep neural network architectures are tested for feature extraction and compared in terms of their ability to detect right whales which can be divided in multiple categories:

- Early architectures: Alexnet, VGG16, VGG19 [48])
- Residual networks: Resnet18, Resnet 50, Resnet101[49]
- Dense networks: Densenet 201[50]
- Dag Networks: Darknet19 and Darknet53 developed as backbone for localization and bounding box estimation frameworks such as YOLO [51,52].
- Efficient and mobile architectures: Nasnet Mobile [49], Mobilenet and MobilenetV2[53], Squeezenet[54], Shufflenet[55], designed for mobile device with less computing power, e.g. Shufflenet reduces computation costs with a 13x speed up while maintaining an equivalent accuracy to Alexnet while SqueezeNet achieves equivalent accuracy as Alexnet with 50 times fewer parameters.
- Large architectures: Nasnet-A [49], InceptionV3 [50], *Inception-ResnetV2 [58], Googlenet [59], Xception [60].*

The original dataset is balanced with, 1160 images, that is two classes of 580 images for each of the two classes. The other class is composed of sea surface and surfacing elements that were visually assigned to the non-whale class (patches of algae, rocks, etc).

In order to make the feature classification more robust by providing more samples for training, the following data augmentation techniques were applied:

1. image scaling within the range [0.5; 1] by steps of 0.05
2. image rotation in the range [-70°; 70°] randomly generated by steps of 1 degree).

This offline data augmentation process produced an artificial dataset of 1,644,060 images out of the original 1160 images. The extracted feature vectors are then randomly split into an augmented training dataset of 986,436 images (60% of the dataset), and a test dataset of 657,624 images (40% of the dataset.

In order to take advantage of the multispectral components, the image bands are filtered by groups and the extracted features are then concatenated as described in figure 2.4. Even though the deep neural networks at stake are originally intended for triband RGB images, it is straightforward to apply the process to a multispectral image by concatenating outputs as described in figure 2.4, with the sole limitation of using band triplets as inputs to the feature extractors.

**Figure 2.4.**
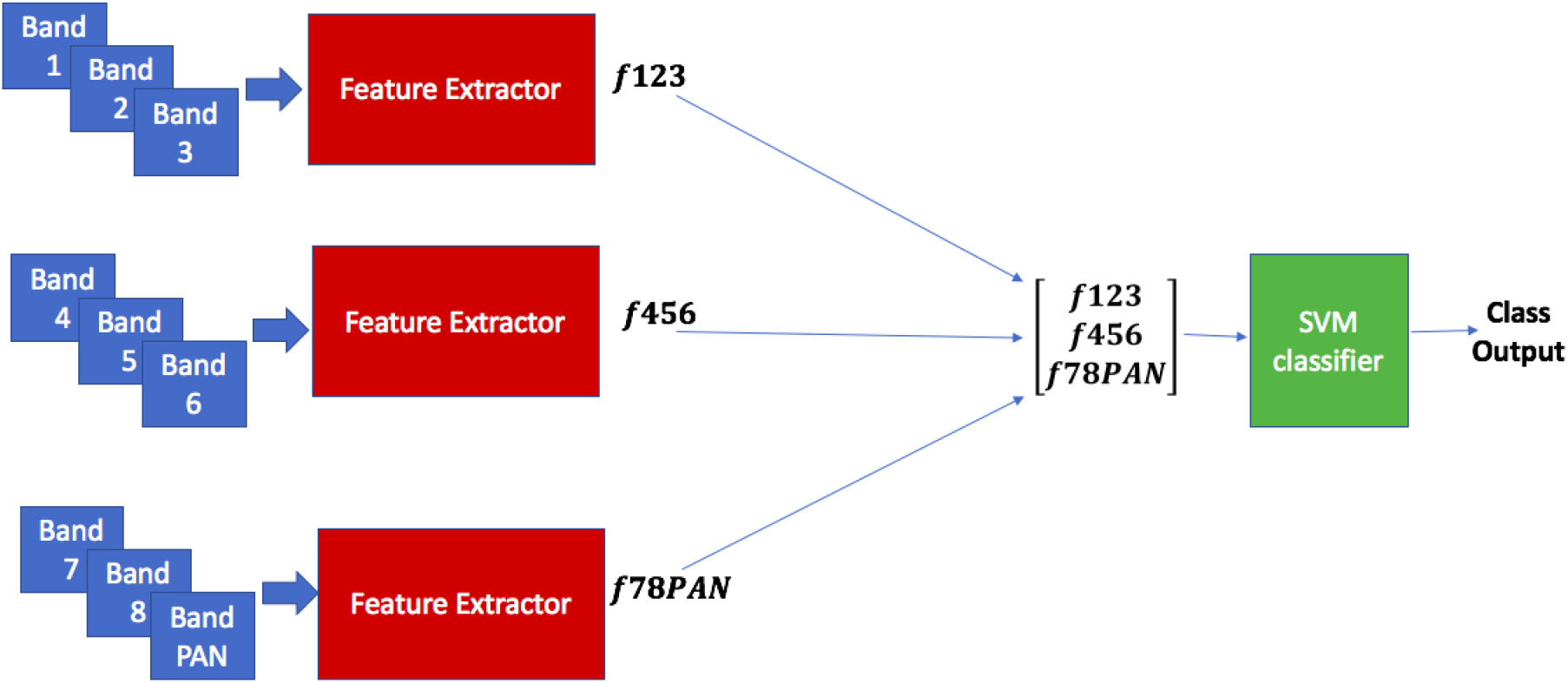
Schematic of feature Extraction and classification with data input V_ALL_.

The following band combinations are tested for detection: V_123_={Coastal(1), Blue(2), Green(3)}, V_RGB_={ Red(5); Green(3)}, Blue(2)}, V_456_ ={ Yellow(4), Red(5), Red-Edge(6)}, V_345_ ={Green(3), Yellow(4), Red(5)}, V_678_ ={Red-Edge(6), NIR1(7), NIR2(8) }, V_678_ ={ Red-Edge(6), NIR1(7), NIR2(8) }, V_1:6_ ={ [Coastal(1), Blue(2), Green(3)], [Yellow(4), Red(5), Red-Edge(6)]},V_1:8_={ [Coastal(1), Blue(2), Green(3)], [Yellow(4), Red(5), Red-Edge(6)], [Red-Edge(6), NIR1(7), NIR2(8)]}, V_ALL_={ [Coastal(1), Blue(2), Green(3)], [Yellow(4), Red(5), Red-Edge(6)],[ NIR1(7), NIR2(8), PAN]}, V_PAN_ ={PAN,PAN,PAN}, V_GEO_ ={ [Blue(2), Green(3), Red(5)], [Green(3), Red(5), NIR1(7)] }. V_PAN_ can be used to describe a panchromatic-only sensor such as Worldview 1 and V_GEO_ describes the spectral content of sensors such as those of Geoeye or Ikonos (see table 2.1).

Deep learning architectures for localization such as Faster-RCNN have been used in previous studies for the detection of marine mammals [61], but the aerial and satellite images used were restricted to the RGB bands, and the small sample support conjoined with a fine-tuning approach typically resulted in a large number of false positives with an F-score of 81% for detection and a 94% accuracy in counting. A benchmark study used non-supervised ad-hoc detection techniques looking into multiple multispectral bands and obtained an accuracy of 84.6% to 89.1% by thresholding the coastal band [14].

#### 2.2.5 Segmentation, region analysis and form analysis

##### 2.2.5.1 Segmentation and region analysis

It is of great interest to be able to extract shape information from the detections of interest. This allows to provide a rough estimate of the dimensions of the whales and to differentiate calves, juveniles and adult specimens. To extract this information automatically, a series of classical morphological operations [62] are used. First, in order to obtain a binary segmentation of the environment, an edge detector with a Prewitt operator for edge detection is used (fig. 2.5a). To better delineate the contour of the whales, the image is then dilated using a circular structural element (fig. 2.5b), filled (fig. 2.5d), smoothed using erosion with a diamond structural element (fig. 2.5e). The final result of the segmentation process is a binary mask (figure 2.5f), from which it is straightforward to extract outlines (fig. 2.5g) and bounding boxes (fig. 2.5g), and finally the properties of the detections of interest can be extracted automatically with region properties algorithms.

**Figure 2.5a to 2.5h:**
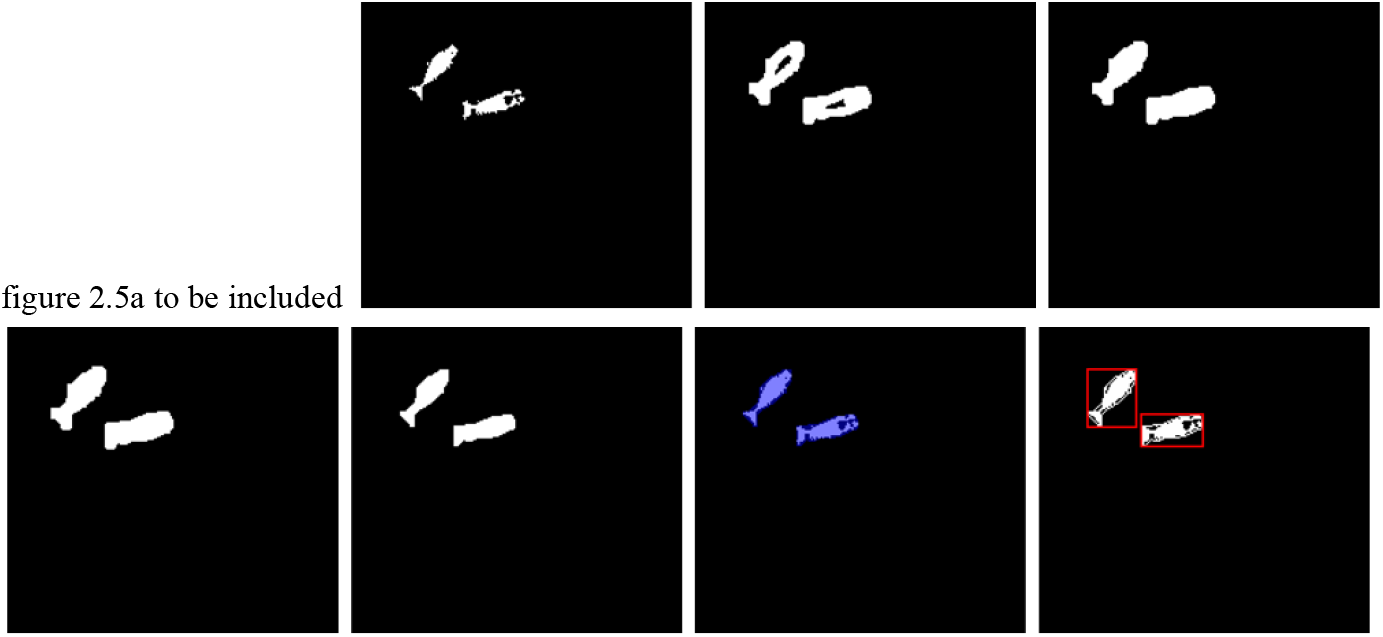
example of morphological operations. Figure 2.5h includes the automatically estimated bounding box around each right whale.

The following properties were focused on:

1. Major Axis length: the estimated length in pixels of the major axis of the ellipse that has the same normalized second central moments as the region.
2. Minor Axis length: the estimated length in pixels of the minor axis of the ellipse that has the same normalized second central moments as the region.
3. Area: the number of whale pixels in the region, returned as a scalar.
4. Eccentricity: the eccentricity of the ellipse that has the same second-moments as the region of whale pixel, that is the ratio of the distance between the foci of the ellipse and its major axis length.
5. Bounding Box Length and Bounding Box width: the bounding box is mathematically the smallest box containing the whale pixels region.
6. Extent: the ratio between the number of whale pixels in the region to pixels in the total bounding box. Computed as the Area divided by the area of the bounding box.
7. Perimeter: the distance in pixels around the boundary of the whale pixel region returned as a scalar.

##### 2.2.5.2 Form analysis

We name ‘typical form’ a right whale observation where the whale can be seen from head through the streamlined body down to the flukes. A typical form does not require fins to be visible. We name atypical form all other right whale observations. This includes multiple whale postures such as spyhopping and tailslapping. This also includes untypical situations such as lunge feeding in which case the body of the whale can be deformed and change orientation. While measurements of typical forms could be used as indicator regarding the whale status (adult, juvenile), measurements of atypical forms are not likely to translate into relevant information.

## 3. Results and Discussion

Using segmented images, one can then extract the following type of statistics can readily be extracted respectively for the water and whale pixels:

1. Visualizations: histograms and scatterplots
2. Numerical estimates: mean value 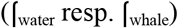, standard deviation 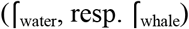, kurtosis (k_water_ resp. k_whale_)*
3. Distribution measurements: the Kullback-Leibler divergence (equation 3.1) is used to measure the similarity between the whale and water probability distributions of the classes of interest derived from their histograms. A large Kullback-Leibler divergence value indicates a larger difference of the probability distributions which as a result makes them more likely to be separable. The overlap in percent between distributions is also estimated from the histograms.
4. In order to account for the variability of information across the multiple available spectral bands, covariance matrices C_water_ and C_whale_ are also extracted.

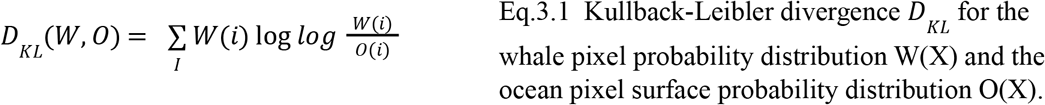

### 3.1 Statistical review of the panchromatic and multispectral bands

The distribution of whale pixels and ocean surface pixels have different sample size which makes it difficult to compare them through raw histograms, instead a probabilistic normalization is applied so that all of the histogram bar heights add to 1 and a uniform bin width is used. We can then estimate the probability density function of the whale pixel W and the probability density function O of the ocean surface and calculate the Kullback-Leibler divergence between the two of them. The larger the divergence between the two distributions, the easier it is to separate the two pixel distributions.

As shown in the series of histograms in figure 3.1(a) to 3.1(i), even though the water and whale pixel distributions are clearly identifiable, there is a significant overlap between them. The panchromatic histogram provides the most overlap which is corroborated by numerical measurements (table 3.1), which indicates that all of the multispectral color bands provide better raw separation capabilities. As can be seen in figures 3.1a to 3.1i, Southern Right Whale pixels are represented in red and typically have lower intensity values whereas water pixels, represented in blue, typically have higher intensity values. We may note that the water pixel distribution has a very regular Gaussian shape, which makes it convenient for the detection of other salient pixels, whereas the whale pixel distribution features an elongated tail which corresponds to lighter pixels such as callosities, white marks on the body, lighter calves or even to rare but not uncommon albinos right whales. The non-normalized mean pixel values for the respective classes are given by 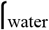 and 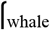 in Table 3.1, whereas normalized pixel values are displayed in the histograms.

**Figure 3.1a to 3.1i.**
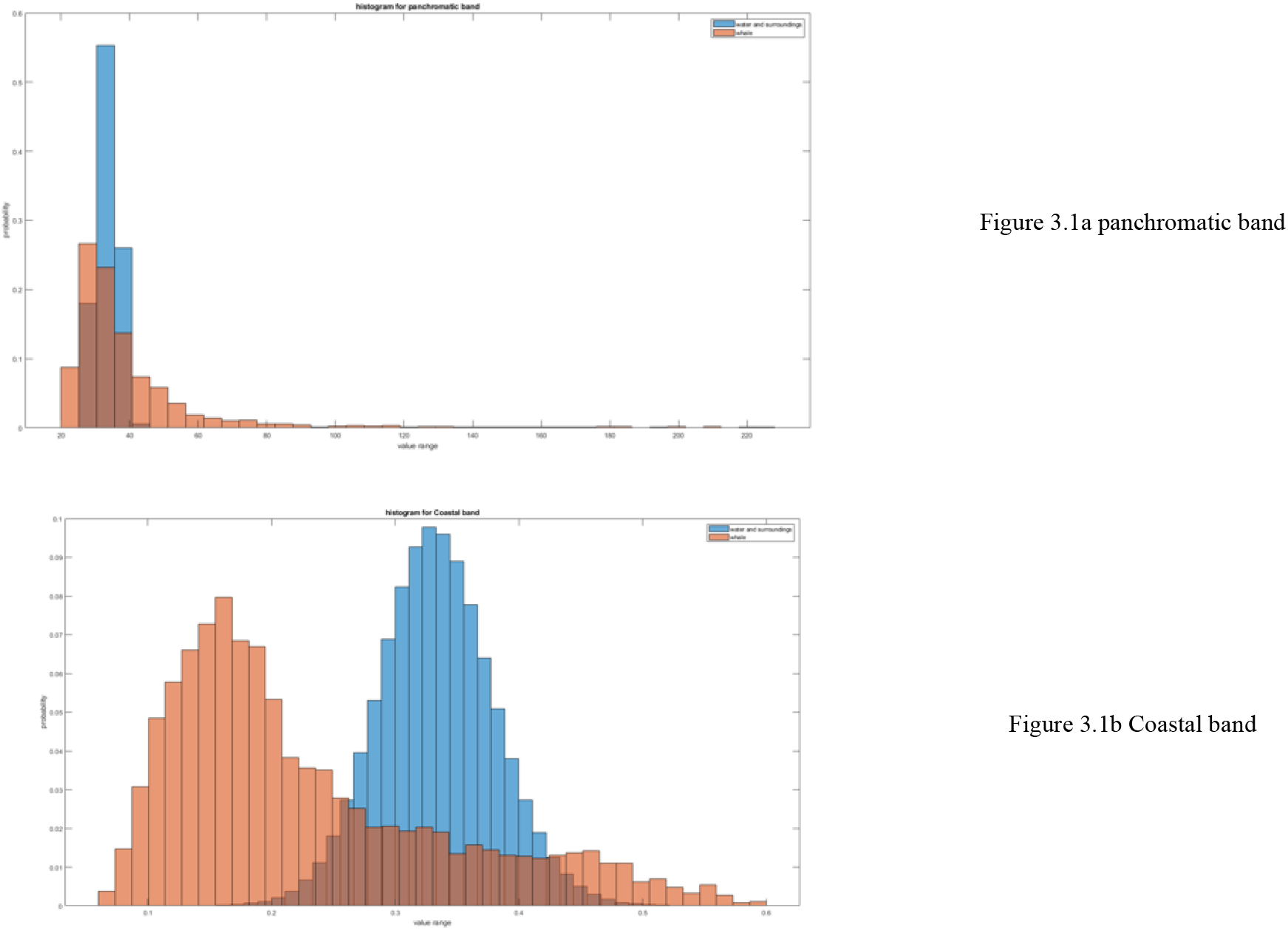
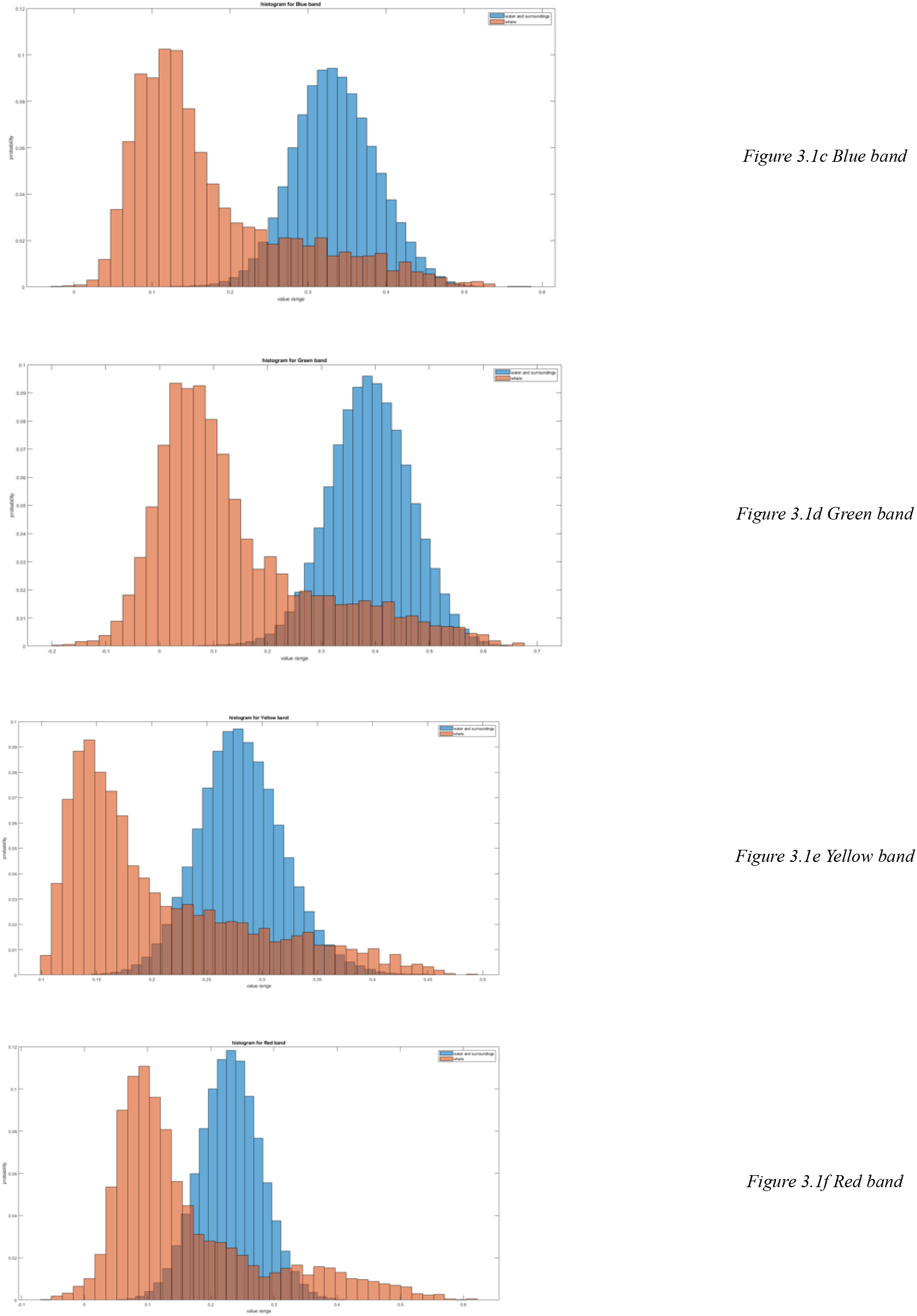
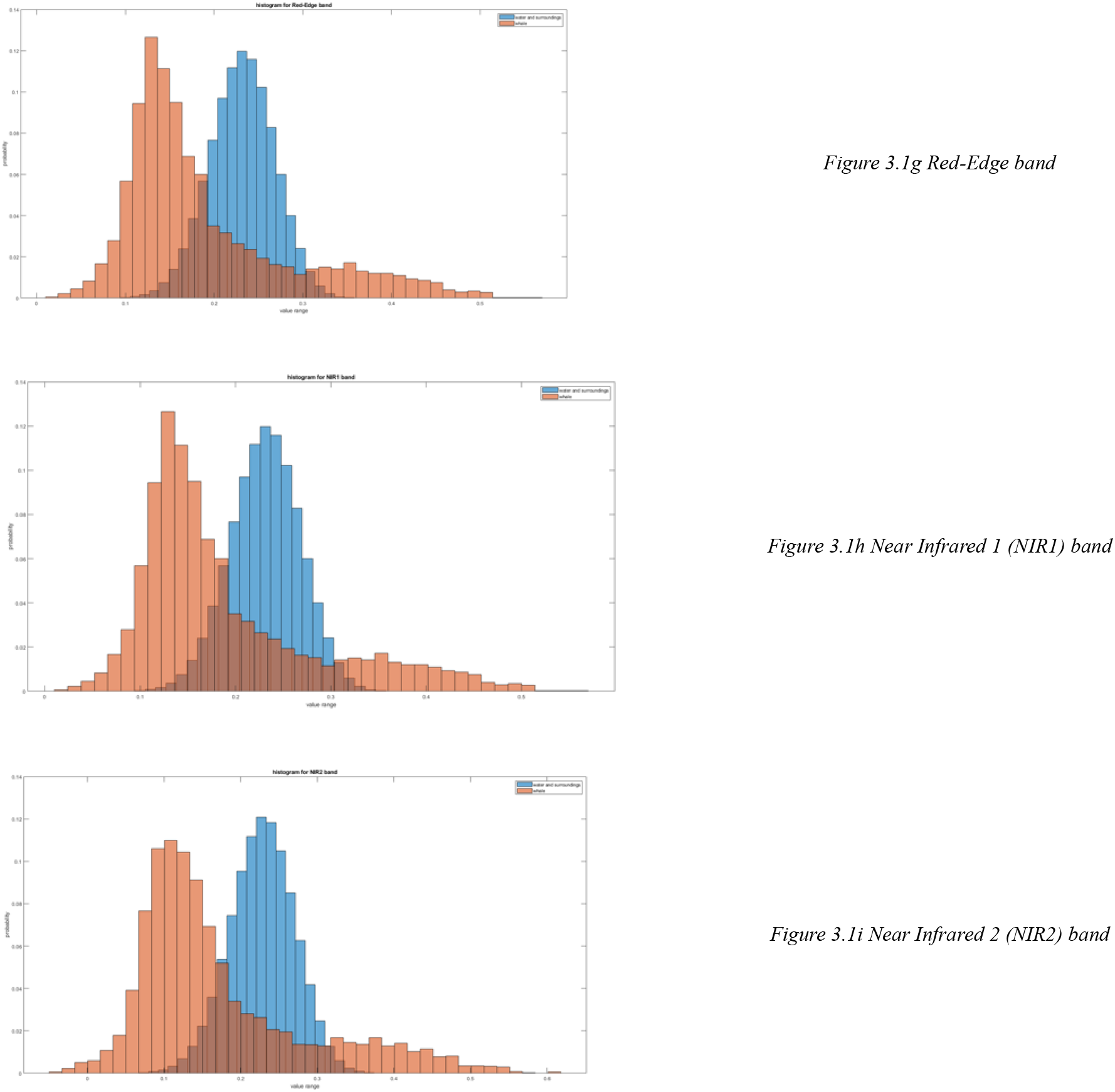
Probability distribution of pixel value per band. The Panchromatic, Coastal, Blue, Green, Yellow, Red, Red-Edge, NIR1 and NIR2 bands are successively represented.

**Table 3.1.** Estimated metrics across panchromatic and multispectral bands for the (non-normalized) whale and the water pixels distributions.

|  | PAN | Coastal | Blue | Green | Yellow | Red | Red-Edge | NiR1 | NIR 2 |
| --- | --- | --- | --- | --- | --- | --- | --- | --- | --- |
| KL | 0.9454 | 1.4257 | 1.7136 | 4.5342 | 6.0703 | 3.2251 | 2.4483 | 2.4483 | 2.9947 |
| f<sub>water</sub> | 30.7997 | 2.9487 | 2.9523 | 3.0473 | 1.6820 | 2.8674 | 2.8853 | 2.8853 | 2.8949 |
| f<sub>whale</sub> | 36.9084 | 2.3921 | 2.2465 | 1.5296 | 2.2354 | 1.5039 | 1.9126 | 1.9126 | 1.6820 |
| f<sub>water</sub> | 3.4217 | 0.0245 | 0.0332 | 0.0693 | 0.0200 | 0.0245 | 0.0141 | 0.0141 | 0.0200 |
| f<sub>whale</sub> | 21.1161 | 0.2280 | 0.2345 | 0.0104 | 0.1058 | 0.2748 | 0.1723 | 0.1723 | 0.2152 |
| Overlap (%) | 22.5 | 10.39 | 5.08 | 5.13 | 5.05 | 5.13 | 5.37 | 5.37 | 5.05 |

Kullback-Leibler divergence (equation 3.1) is used to estimate the separation between both distributions. It is observed that the maximum separation is achieved for Green, Yellow and Red bands, while the Pan, Coastal, and Blue bands provide the least separation between whale and water pixels. An overlap value between distributions in percentage was also estimated where the least overlap was found for NIR2, blue and Yellow band, whereas the most overlap was found for the panchromatic and Coastal bands. In both classes non-normalized variance estimates are found to be greater for the panchromatic band, which highlights the difficulty of performing intensity-based separation using that single band. Across colors, variance is typically greater for whale pixels which highlights potential limitations for the color-based detection. Lesser variance is found across the Blue, Coastal and Yellow bands for both distributions.

Finally, kurtosis μ_4_ indicates that the water distribution is concentrated around, particularly in the Green band but also significantly across the Red-Edge, NIR1 and NIR2, indicating that those bands ae more typical of water reflectance, this is consistent with the rapid absorption of near-infrared light in water whereas the presence of whales causes reflections of those components of the spectrum which leads to very distinct distributions.

As could be expected, the panchromatic band does not separate whale pixels from the surrounding ocean pixels, mean values are relatively close and Kullback-Leibler divergence is close to 1, indicating probability density functions that are very similar and with a significant overlap of 22.5%. For the pansharpened multispectral components, the highest separation as measured by Kullback-Leibler divergence is observed in the Green, Yellow and Red Bands (GYR) while the worst separation is observed in the coastal, blue and NIR bands (CBN1). This is confirmed by the overlap values of the estimated probability density functions even though differences are not as striking. It is remarkable that the Red-Edge and NIR1 bands produce identical results on all metrics, which could indicate that one of this bands does not bring additional information for detection. In terms of visualization a triband RGB image and a triband GYR image, which mathematically provides a better separation of whale pixels and water surface pixels, are shown in figures 3.2(a) and 3.2(b). This suggests that using the GYR image as opposed to the standard RGB image might help visual assessment and that those bands should be given a higher weight for automated detection systems which will be presented in section 3.3. The Green band is of particular interest here as it has the highest mean value for the water pixels (mu0) and the lowest mean value for the whale pixels (mu1) indicating the center of distributions to be farther apart. As can be seen from the histograms in figure 3.1, while the separation between the mean values of the distribution is relatively large, there remains large overlapping region across all spectral bands. These overlapping areas can reasonably be expected to increase rapidly as the depth of the southern right whales in the water column increases.

**figure 3.2a (left) and 3.2b (right).** RGB and GYR representation of a scene containing 7 closely located southern right whales.

### 3.2 Covariance matrices

The sample covariance matrices C_whale_ and C_water_ is estimated for both the non-normalized whale and water pixels. The observed clear difference in range highlights the stronger covariance across bands in the whale distribution than across the water surface distribution. In that sense, the water surface could be understood as a form of correlated noise with a signal to noise ratio varying across bands from 10.9dB (in the NIR1 and Red-Edge Band where water pixels are less correlated) to 7.23dB (in the Yellow band). These significant differences in covariance makes it a strong statistical separator which can potentially be used for robust detection. Particularly when few samples are available the covariance matrix information is an asset which can be directly used by researchers in order to perform detection [63, 64]. The extracted covariance matrices also show us a peculiar characteristic: the Red-Edge band and first Near-Infrared band appear nearly identical for the chosen precision (columns 6 and 7 in covariance matrices and table 3.1). This is not surprising as those bands are most frequently used for vegetation index measurements [65,66] to assess the health status of plants and this suggests that the use of both bands simultaneously does not yield significant discrimination capabilities for the current problem.

|  |  |  |  |  |  |  |  |
| --- | --- | --- | --- | --- | --- | --- | --- |
| $C_{\text{whale}}=$ | | | | | | | |
| 0.0520 | 0.0532 | 0.1363 | 0.0214 | 0.0622 | 0.0389 | 0.0389 | 0.0480 |
| 0.0532 | 0.0550 | 0.1407 | 0.0217 | 0.0640 | 0.0397 | 0.0397 | 0.0493 |
| 0.1363 | 0.1407 | 0.3726 | 0.0507 | 0.1663 | 0.1031 | 0.1031 | 0.1303 |
| 0.0214 | 0.0217 | 0.0507 | 0.0112 | 0.0246 | 0.0158 | 0.0158 | 0.0182 |
| 0.0622 | 0.0640 | 0.1663 | 0.0246 | 0.0755 | 0.0470 | 0.0470 | 0.0586 |
| 0.0389 | 0.0397 | 0.1031 | 0.0158 | 0.0470 | 0.0297 | 0.0297 | 0.0367 |
| 0.0389 | 0.0397 | 0.1031 | 0.0158 | 0.0470 | 0.0297 | 0.0297 | 0.0367 |
| 0.0480 | 0.0493 | 0.1303 | 0.0182 | 0.0586 | 0.0367 | 0.0367 | 0.0463 |
| $C_{\text{water}}=$ | | | | | | | |
| 0.0006 | 0.0008 | 0.0016 | 0.0004 | 0.0006 | 0.0003 | 0.0003 | 0.0004 |
| 0.0008 | 0.0011 | 0.0023 | 0.0006 | 0.0008 | 0.0004 | 0.0004 | 0.0005 |
| 0.0016 | 0.0023 | 0.0048 | 0.0012 | 0.0017 | 0.0009 | 0.0009 | 0.0012 |
| 0.0004 | 0.0006 | 0.0012 | 0.0004 | 0.0004 | 0.0002 | 0.0002 | 0.0003 |
| 0.0006 | 0.0008 | 0.0017 | 0.0004 | 0.0006 | 0.0003 | 0.0003 | 0.0005 |
| 0.0003 | 0.0004 | 0.0009 | 0.0002 | 0.0003 | 0.0002 | 0.0002 | 0.0003 |
| 0.0003 | 0.0004 | 0.0009 | 0.0002 | 0.0003 | 0.0002 | 0.0002 | 0.0003 |
| 0.0004 | 0.0005 | 0.0012 | 0.0003 | 0.0005 | 0.0003 | 0.0003 | 0.0004 |

### 3.2 Measurements and observation of typical, atypical forms and mother and calf pairs

Extracted whale measurements are presented in Table 3.2. for typical and atypical forms. While dimensions are influenced by the particular situation of the image (e.g. nadir or off-axis angle of the satellite’s sensor), such automatically extracted measurements can provide important information regarding the difficulty of the task at hand. They indicate that a large diversity of situations can be observed that will influence the ability of an automated system to perform detection. Typical forms, are generally whales located on or near the ocean surface which makes them ideal for human or automated recognition. Their dimensions as evidenced by the major axis and minor axis measurements are larger on average and have less variance, while atypical forms are typically smaller and with greater variance. The study area was purposely selected for its shallow waters and excellent visibility conditions so that typical forms dominate our observations, however, in more adverse situations, atypical forms can be expected to be dominant. Due to their smaller size and larger variance, they can be expected to make automated detection more difficult. They represent a greater variety of situations with whales located deeper in the water column or in a larger variety of postures (tailslapping, spyhopping, etc). In that sense, a best performing algorithm should be able to operate correctly with the reduced pixel support of atypical forms. The typical/atypical distinction is therefore useful to know what to expect in more adverse situations (higher sea state, clouds, fog, etc). They also represent situations where salient features that are typical of right whales and cetaceans in general (such as their tail) cannot be observed. Algorithms trained only with typical forms could therefore fail to recognize atypical forms. As the ability to recognize whale and their features decreases with resolution, atypical form are also an indication of how the algorithm can perform for sensors with a lower resolution.

**figure 3.3.** Example of typical forms observed in World View 3.

**figure 3.4.** Example of atypical forms observed in World View 3.

**figure 3.5.** Example of mother and calf pairs observed in World View 3.

**Table 3.2:** Extracted form measurements for typical and atypical forms.

| Measurement | Typical (Essential) forms |  | Atypical forms |  |
| --- | --- | --- | --- | --- |
|  | Mean | Standard deviation | Mean | Standard deviation |
| Major axis | 45.0 | 13.9 | 32.9 | 21.4 |
| Minor Axis | 19.2 | 6.2 | 15.1 | 9.6 |
| Area | 668.1 | 259.4 | 484.4 | 374.2 |
| Bounding Box Width | 35.7 | 13.6 | 26.3 | 18.1 |
| Bounding Box Height | 34.6 | 12.7 | 26.1 | 18.4 |
| Eccentricity | 0.9 | 0.04 | 0.8 | 0.23 |
| Extent | 0.5 | 0.1 | 0.6 | 0.2 |
| Perimeter | 119.7 | 28.2 | 91.7 | 62.8 |

### 3.3 Result of Feature Extraction and detection

For a proper evaluation of the multiple pretrained models, a 5-fold cross-validation was employed and data was randomly subdivided in a 60% training subset and a 40% testing subset, results are then expressed as average accuracy, average sensitivity, average specificity and average precision. Feature extraction followed by classification achieved consistently high accuracy results over the dataset with important variations depending on the trained neural network used as backbone. The 5 best average network performances and the 5 worst average network performances are presented respectively in tables 3.3 and 3.4 for rapid examination, while overall results are presented in table 3.5. While overall results seem excellent, perfect performance is only achieved in two configurations with the Nasnet-A network and the Inceptionv3 network, and both those results are only achieved with the multispectral bands of V345. This highlights the importance of band and architecture selection for detection and classification. While seemingly excellent results could also be achieved with Shufflenet, nasnetmobile and squeezenet, the appearance of false positives with a false positive rate of 5.04% could result in a complex review process in the context of a big data problem. A single satellite image typically involves hundreds of millions of pixels. The explanation of the excellence of certain networks and the poorer performance of others is not straightforward but we may notice that 4 of the best performing architectures Nasnet large, Shuf?lenet, Squeezenet and nasnet mobile are lightweight convolutional neural network architectures, with an improved computational efficiency meant for mobile devices. Squeezenet theoretically has the same theoretical accuracy level as AlexNet on the IMAGENET dataset with a smaller model size and 50 times fewer parameters [47], which, on top of its performance, is a major asset for the processing of large datasets. Here squeezenet performed significantly better than its Alexnet counterpart which at best achieved a 95.12% accuracy (V_GEO_), it still provided twice the best response for some suboptimal band selections (V_456_ and V_678_). NASNet Large was also at the time of its publication a step forward in performance with a 28% reduction in computational demand through the use of a simpler model [56]. Inception v3 was itself a modification of earlier Inception CNNS with the clear objective of reducing memory and computational demands [57]. It is remarkable that the five best networks do not produce any false negatives. Smaller networks like Shufflenet, Nasnetmobile and squeezenet performed high and maintained a 100% sensitivity whereas for all other responses the degradation of performance occurs mostly through a decrease of sensitivity, i.e. through an increased number of false negatives. Such false negatives are critical and should be avoided for detection systems that deal with animals with an endangered status for which each individual detection matters.

**Table 3.3.** 5 best network performances on the test data set.

|  | Nasnet | INCEPTIONV3 | Shufflenet | Nasnetmobile | SQUEEZENET |
| --- | --- | --- | --- | --- | --- |
| Average Accuracy | 100% | 100% | 98% | 98% | 98% |
| Average Specificity | 100% | 100% | 95% | 95% | 95% |
| Average Sensitivity | 100% | 100% | 100% | 100% | 100% |
| Average Precision | 100% | 100% | 95% | 95% | 95% |
| Average True Positives | 116 | 116 | 113 | 113 | 113 |
| Average True Negatives | 116 | 116 | 113 | 113 | 113 |
| Average False Positives | 0 | 0 | 6 | 6 | 6 |
| Average False Negatives | 0 | 0 | 0 | 0 | 0 |

**Table 3.4.** 5 worst network performances on the test data set.

|  | Googlenet | Resnet18 | Mobilenetv2 | Shufflenet | Resnet101 |
| --- | --- | --- | --- | --- | --- |
| Average Accuracy | 80% | 80% | 83% | 83% | 85% |
| Average Specificity | 95% | 90% | 90% | 95% | 95% |
| Average Sensitivity | 65% | 70% | 75% | 70% | 75% |
| Average Precision | 92% | 88% | 88% | 93% | 94% |
| Average True Positives | 73 | 79 | 85 | 79 | 90 |
| Average True Negatives | 113 | 108 | 108 | 113 | 113 |
| Average False Positives | 6 | 11 | 11 | 6 | 6 |
| Average False Negatives | 40 | 34 | 28 | 34 | 23 |

|  |  | 1 | 2 | 3 | 4 | 5 | 6 | 7 | 8 |
| --- | --- | --- | --- | --- | --- | --- | --- | --- | --- |
|  |  | Best accuracy | Networks for Best accuracy | Worst accuracy | Networks for worst accuracy | Mean and Standard Deviation of accuracy | Mean and Standard Deviation of precision | Mean and Standard Deviation of sensitivity | Mean and Standard Deviation of specificity |
| 1 | V<sub>123</sub> | 97.56% | Shufflenet<br>Squeezenet | 87.80% | Densenet201<br>Nasnet Large<br>Googlenet<br>Resnet18<br>Resnet101<br>Vgg19<br>Mobilenetv2 | 91.3/<br>3.4 | 93.5/<br>3 | 88.4/<br>7.5 | 93.9/<br>3.2 |
| 3 | V<sub>RGB</sub> | 92.68 | Darknet19 | 82.93% | Squeezenet | 88.96/ | 92.37 | 86.07/ | 93.24 |
|  |  |  | Darknet53 |  |  | 2.62 | 3.38 | 4.87 | 3.29 |
| 4 | V<sub>456</sub> | 92.68% | Darknet19<br>Resnet50<br>Alexnet | 80.49% | Resnet18 | 88.5/<br>3.3 | 94.43/<br>4.63 | 82.06/<br>5.6 | 95.24/<br>3.27 |
| 5 | V<sub>345</sub> | 100% | Nasnetlarge,<br>Inceptionv3 | 90.24% | Vgg16 | 96.34/<br>2.40 | 94.62/<br>4.21 | 98.33/<br>2.97 | 94.18/<br>4.49 |
| 6 | V<sub>678</sub> | 90.24% | Alexnet | **80.49%** | Googlenet<br>Resnet18 | 86.87<br>/3.49 | 93.16<br>/1.97 | 77.5<br>/3.93 | 95.03<br>/2.26 |
| 7 | V<sub>1:8</sub> | 95.12% | Nasnet large | 85.37% | mobilenetV2<br>Vgg19 | 89.16/<br>2.67 | 95.49/<br>1.92 | 83.42/ 5.54 | 94.30<br>1.99 |
| 8 | V<sub>1:6</sub> | 97.56% | Squeezenet | 85.37% | mobilenetV2<br>densenet201 | 89.24/<br>2.29 | 93.82/<br>3.62 | 85.56/<br>6.84 | 94.49/<br>3.97 |
| 9 | V<sub>GEO</sub> | 97.56% | Darknet19<br>Resnet50<br>Googlenet | 82.93% | densenet201<br>Vgg19 | 90.38/<br>4.77 | 94.27<br>4.67 | 85.56/<br>7.65 | 94.97<br>4.16 |
| 10 | V<sub>PAN</sub> | 95.12% | Shufflenet | 75.61% | Resnet50 | 85.64/<br>5.79 | **92.73/**<br>**7.26** | 76.67/<br>7.28 | 94.18/<br>5.79 |
| 11 | V<sub>ALL</sub> | 92.68% | darknet19'<br>'resnet101' | 82.93% | 'inceptionv3'<br>'vgg16' | 88.35<br>2.84 | **95.12**<br>**4.80** | 80.56<br>5.91 | 95.77<br>4.29 |
| 12 | V<sub>OCI-NI</sub><br><sub>R1</sub> | 95.12% | Resnet50 | 82.93% | nasnetmobile | 89.47/3.15 | **93.08/4.81** | 85/5.53 | 93.73/0.045 |
| 13 | V<sub>OCI-NI</sub><br><sub>R2</sub> | 97.56% | Resnet50 | 75.61% | alexnet | 88.58/5.33 | **94.44/7.06** | 81.84/7.49 | 94.99/7.00 |

As could be expected from the analysis of the probability distribution per multispectral band (section 3.1), V_345_, a vector composed of all bands with the largest Kullback-Leibler divergence provided the best results (100% average accuracy) and its worst performance, achieved with Vgg16, is still relatively high (90.24% average accuracy). V_678_ obtained the smallest best average accuracy (C_6,1_), even surpassed by the panchromatic band (C_10,1_) even though the latter got the smallest worst average accuracy (C_10,3_) and performed worse on average across networks (C_10,5_). The performance of V_678_ is not entirely surprising since the Red-Edge and Near Infrared 1 bands provide the same response in all metrics of table 3.1, therefore indicating that one of those bands does not apparently bring much information for classification of this dataset. The typical role of the red-edge and near infrared bands that has proven useful in agriculture to classify and assess the health status of vegetation [65,66] does not seem to yield discrimination benefits for the current task. It could be inferred that the rapid absorption of NIR underwater may bring randomness to the data. For whales located deeper in the water column where the near-infrared light has been rapidly absorbed, those components will therefore not yield strong discrimination ability between whale and water.

More complex networks such as Resnet, ExceptionV3 and Densenet who typically have better results on large standards datasets such as Imagenet, do not perform better for feature extraction and classification of the dataset. Larger amounts of data for training could change these dynamics and the results highlighting the best performing network should be updated as the number of detection keeps increasing, before eventually turning to a full network training when enough data is gathered. It should also be noted be that at equal performance smaller architecture should be favored so as to reduce complexity and computation time.

#### Conclusions and Future Work

A key result has been to extract significant statistics that can be used to develop a wide range of approaches for detection (from radial basis function to GMM) which do not require a large sample support contrary to deep-learning methods. A deep-learning-based architecture was also developed that uses multispectral and spatial information contained in all the multispectral bands and performs at 100% accuracy, a result which sketches an interesting alternative to fine-tuning a deep neural network with a small sample support which generally results in higher False Positive and False Negative Rates [61]. Furthermore, no such approach has yet been proposed for whales beyond the RGB channels which were demonstrated in the statistical analysis to yield suboptimal results. The proposed methods by avoiding false positives and negatives allows for a rapid review of satellite data and a rapid increase of the number of detections.

Overall performance is relatively good, with the two best networks committing no mistake for the detection/classification of southern right whales instances. Even whales located deeper in the water column, representing fainter signals, are detected. In the context of big spatial data, where even low false positive rates produce a number of false positives that can rapidly become intractable for human review, those results are extremely encouraging. While performance can be expected to deteriorate under more adverse conditions with wind, waves and clutter and possibly with more instances that could be misclassified as whales, the detector and extracted statistical information already provide a basis to rapidly increase the number of detections which will in turn allow to build and train increasingly robust detectors. Future work will involve retraining the algorithm and increasing its performance in a variety of conditions, the inclusion of a variety of other events that could potentially occur (seabirds, waves, ship, fishing nets, rocks, other marine mammals, etc.). A future paper will focus on the deployment of the upgraded detector in other areas particularly for the more endangered populations of North Pacific Right Whales and North Atlantic Right whales. The algorithm could furthermore be deployed on larger datasets in South Africa to study yet unexplained distributional shifts.

## Acknowledgments

We thank Dr. David Weller for his kind sponsorship and helpful review of this study.

We thank the Eppley Foundation For Research for their kind support of the Wildlife and Marine Mammal Spatial Observatory project.

## Notes

### Competing Interest Statement

The authors have declared no competing interest.

